# A severe form of maternal separation in rat consisting of nineteen-day 6-hour daily separation at unpredictable times minimally affects behavior across the lifespan

**DOI:** 10.1101/2021.06.07.447289

**Authors:** Tatyana B. Behring, Margaret H. Kyle, Maha Hussain, Jack Y. Zhang, Alessia Manganaro, Jasmine H. Kaidbey, Robert J. Ludwig, Michael M. Myers, Martha G. Welch, Dani Dumitriu

## Abstract

Maternal separation (MS), a type of early life stress, has been associated with adverse socioemotional and behavioral outcomes throughout the lifespan across multiple species. Comprehensive longitudinal biobehavioral characterization of MS in rats is sparse and conflicting, warranting more studies. We conducted an MS paradigm involving 6-hour daily separation at unpredictable start times from P2 to P21. We hypothesized this severe form of MS would lead to developmentally emerging maladaptive biobehavioral consequences from juvenile through adult periods compared to Controls (C), especially in social behaviors. We tested: (1) own dam odor preference shortly after weaning; (2) juvenile and adult anxiety-like, sociability, and play behaviors using the light-dark test, three-chambered social interaction test, and video-coded juvenile play behavior; and (3) adult coping behaviors and neuroendocrine response using the forced swim test and blood corticosterone responses. Our results show minimal effects on biobehavioral outcomes across the lifespan. Recently weaned MS male rats had a stronger preference for their dam’s odor. Juvenile MS females spent more time in rough-and-tumble play than C female rats. No differences in sociability were found in the juvenile or adult periods. MS promoted a decrease in anxiety-like behavior that persisted from juvenile to adult periods. Finally, MS led to deficits in coping behavior in male adults, but basal and reactive corticosterone levels were unaltered by MS. More studies are needed to validate our surprising findings and probe the neural mechanisms underlying some of the observed protective effects.

**Significance Statement:** Maternal separation (MS), a type of early life stress, has been shown to lead to short and long-term adverse socioemotional consequences in humans and biobehavioral outcomes in rodents. Available data on MS exposed rats are sparse, conflicting, and often lack longitudinal outcomes. We designed a 6-hour unpredictable daily MS paradigm from postnatal day (P2) to P21 to severely disrupt normal dam-pup interactions. Our results show MS leads to increased preference for own dam odor, persisting reductions in anxiety-like behaviors from juvenile to adulthood, and no effects on sociability. Adverse effects on coping behaviors were uniquely identified in males. The present study warrants reevaluation of the central dogma that MS leads to maladaptive biobehavioral outcomes.

## Introduction

Early postnatal development is important in determining future socioemotional outcomes. In this period, infants are sensitive to the “loss” of- the mother (Bowlby, 1970, 1982; Hofer, 1996; Welch, 2016; Ludwig and Welch, 2019), and maternal deprivation is broadly classified as an early life stress (Molet et al., 2014; Welch, 2016). Maternal separation (MS) and/or deprivation in humans is associated with adverse socioemotional outcomes in childhood (Kaler and Freeman, 1994; Chisholm, 1998; Chugani et al., 2001), adolescence (Kreppner et al., 2007; Beverly et al., 2008; Stevens et al., 2008; Shin et al., 2016; Pesarico et al., 2017) and adulthood (Mozaffari, 2018). Parallel findings have been reported in rodents (Kalinichev et al., 2002; Romeo et al., 2003). Given MS-mediated adverse outcomes are observed across the lifespan in both humans and rats, this model is critical for identifying mechanisms of early life stress and possible therapeutic strategies for treatment and/or prevention.

The MS paradigm is typically comprised of daily 3-hour separations, but longer separations between rodent dams and pups during the first two weeks of life are also used (Baudin et al., 2012; Xue et al., 2013; Zhang et al., 2013; Lundberg et al., 2017b). The dam’s care for her pups has been shown to mediate many physiological processes such as feeding, nipple attachment, and heart rate (Hofer, 1983; Myers et al., 1989; Kentrop et al., 2018). The disruption of normal interactions by MS and its acute negative effects on these physiological processes have been studied for many years (Hofer and Weiner, 1971; Hofer, 1975; Ranger et al., 2021). Longer-term adverse effects have been associated with impaired social functioning (Kambali et al., 2019), but more frequently with increases in stress responses, anxiety-, and depressive-like behaviors in both dams and pups, as well as increases in corticosterone levels (Stanton et al., 1988) and altered HPA activation in MS pups (Liu et al., 1997; Caldji et al., 2000; Kalinichev et al., 2002; Maniam and Morris, 2010). In contrast, other studies have demonstrated that MS can promote resilience or have few long-term consequences (Lundberg et al., 2017a; Kambali et al., 2019). These conflicting findings and the lack of emphasis on social outcomes warrants further investigation of the paradigm (Lehmann et al., 2000).

Previously, our group demonstrated altered patterns of ultrasonic vocalizations (Kaidbey et al., 2019) and blunted pup fronto-cortical activity during normal in-cage dam-pup interactions (Ranger et al., 2021) in a 3-hour, 10-day MS rat model. In the current study, we sought to establish the effects of a severe form of MS on long-term biobehavioral outcomes in rats. We used a 6-hour daily MS paradigm with an unpredictable start time and isolation from both dam and littermates from postnatal day (P) 2 through P21 in rats (D’Agata et al., 2017). We compared MS-reared animals to standard animal facility reared controls (C) to test for effects of MS biobehavioral outcomes across the lifespan. To assess this, we examined (1) own dam odor preference shortly after weaning; (2) juvenile and adult anxiety-like, sociability, and play behaviors using the light-dark (LD) test, three-chambered social interaction test (SIT, and video-recorded juvenile play behavior; and (3) adult coping behaviors and neuroendocrine response using the forced swim test (FST) and blood corticosterone responses. To the best of our knowledge, this study represents the first rigorous approach to elucidating the effects of early life stress on long-term biobehavioral consequences in a severe form of rat MS.

## Methods

### Subjects

Sixteen timed-pregnant Sprague Dawley dams from Charles River (Wilmington, MA) were received on E17 and housed in the New York State Psychiatric Institute rat colony facility. Animals were kept in a 12-hour dark/light cycles with lights on at 7 am in a temperature (21°C) and humidity controlled (40%) room, with food and water available ad libitum and routine husbandry provided. The dams were checked daily to determine the precise day of birth (postnatal day 0, P0). On P2, litters were culled to 8 with a 1:1 male: female ratio when possible. All procedures were conducted strictly following animal protocols approved by Columbia University Irving Medical Centre and New York State Psychiatric Institute’s Institutional Animal Care and Use Committees.

A total of 104 (1:1 male:female) of 128 rats were selected randomly to be used in the study. All litters (*n*=16 [control=8, maternally separated=8]) were weaned on P21 and transferred to a new cage with 2 rats per cage. A smaller portion of rats (*n*=40 [C=20, MS=20, 1:1 male:female]) was randomly assigned to a cross-housed condition in which they were housed with rats reared differently (control with maternally separated offspring). The rest of the rats were housed with siblings from the same litter (n=64 (C=32, MS=32 [1:1 male:female])). Of these rats, 24 (C=12, MS=12,1:1 male:female) were randomly assigned to a non-tested group, which would not undergo juvenile testing. A flow chart of the study procedures can be found in Figure 1 as well as the weights of the animals.

**Fig. 1.**
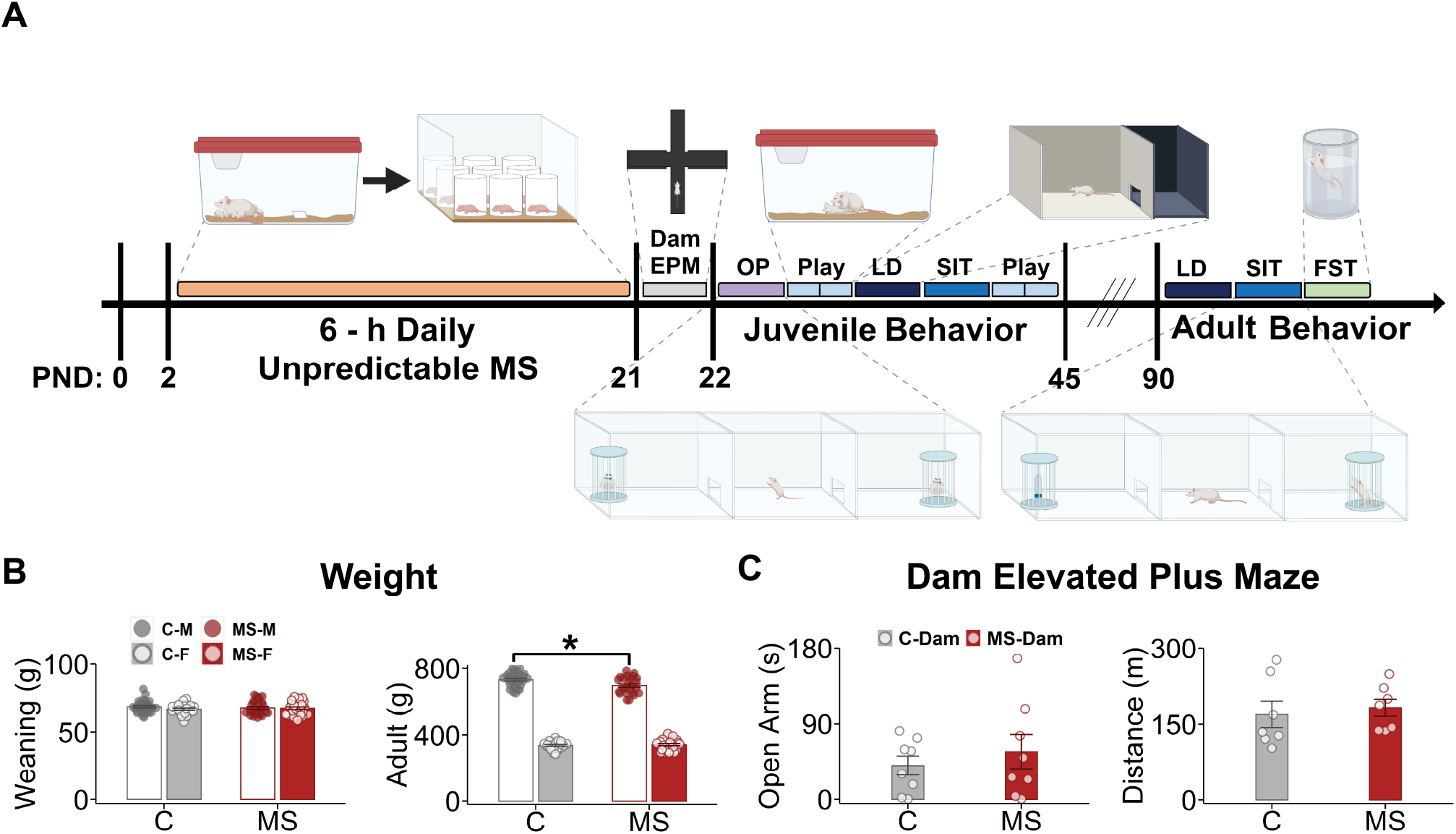
Figure 1. A severe maternal separation (MS) model designed to induce maximal biobehavioral consequences throughout the lifespan. **(A)** Pup births were monitored to establish P0. The 6-h daily unpredictable MS began on P2 and lasted until P21. Pups were separated from their dam and littermates and placed in cylinders for the duration of the 6-hour period. On P21, pups were weaned and dam anxiety was tested on the elevated plus maze (EPM) test. Starting on P22 juvenile behavioral testing began and included maternal odor preference (OP), light-dark box test (LD), social interaction test (SIT) and repeated testing of play behavior. Starting on P90, adult behavioral testing began and included LD, SIT, and forced swim test (FST). **(B)** Weaning (left) and adult (right) weights. No weight effects were observed at weaning (Rearing: *F*_1,99_ = 0.013, *p* = 0.91, Sex: *F*_1,99_ = 1.10, *p* = 0.30, Rearing × Sex: *F*_1,99_ = 0.68, *p* = 0.41); As expected, a sex effect was observed in adult rats, with males weighting significantly more than females (Sex: *F*_1,98_ = 2233, *p* = 2.92 × 10^-69^). Interestingly, while no rearing effect was observed, a significant rearing by sex effect was seen, with MS males, but not females, weighting slightly less (Rearing: *F*_1,98_ = 3.63, *p* = 0.60, Rearing × Sex *F*_1,98_ = 5.82, *p* = 0.018, Tukey’s, *p* = 0.012). **(C)** No significant differences were observed in EPM open arm time (Open Arm (s), *t*(10.9) = −0.708, *p* = 0.49) (left) or total distance travelled (Distance (m), *t*(10.2) = −0.416, *p* = 0.69) (right) in MS versus C dams immediately post-weaning. Bars represent means ± SEM. \**p* < 0.05 on Tukey’s Test.https://www.overleaf.com/project/60bbd38d4716625e076a5b9e

### Maternal separation procedure

From P2 to P21, the maternal separation pups (MS group, 8 litters, 64 pups) were each isolated in cylindrical PVC partitions inside a new cage with fresh bedding (one pup per cylinder, 8 cylinders per cage, Fig. 1A), as previously described in Kaidbey (Kaidbey et al., 2019) and Ranger (Ranger et al., 2021). Pups were separated from their mothers and siblings at randomly selected starting times that were 1.5 hour apart during the light phase of the cycle between 7:30 AM and 12:30 PM (i.e., 7:30, 9:00, 10:30, 11:00, 12:30) for 6 hours a day. The dams were also placed into new cages with food and water available ad-libidum. After the separation period the pups were returned to their home-cage followed by the dam. Pups in the control group remained with their mother until weaning at P21, except for handling once a week for routine husbandry. Pup weight was measured on P21 and P85.

### Testing Procedures

Pups were weaned at P21. Half were co-housed with one same-sex sibling (sibling-housed group) and half with one individual from another litter (cross-housed group). MS animals were co-housed with control animals in the latter situation. Of the 128 pups that were weaned, 104 of those were randomly chosen to be tested. Ultimately, 80 rats from 40 pairs were selected randomly for juvenile behavioral testing and were a part of the “tested group”, while 104 completed adult behavioral testing and included both “tested” and “untested” animals. The “untested” rats did not undergo LD testing during the juvenile period, but did undergo play behavioral testing, since this was minimally stressful and involved their own cage-mates.

### Maternal odor preference test (OP)

The OP test used a 3-chambered box with dimensions 42 × 62 × 30 cm^3^ (custom made, Curbell Plastics, Orchard Park, NY, USA), and cylindrical interactors, with outer diameters of 10 cm and heights of 25cm. The 2 peripheral chambers were separated from the middle section with boards that had 10 cm wide slots for rat movement between chambers. Dam odor was allowed to accumulate in the cage for the first 24 hours of after weaning and was placed into satchels that were placed into the cylindrical interactors. The location of the unfamiliar dam odor was alternated with each experiment to avoid place-preference. Animals were initially placed in the apparatus for a 5-minute habituation period without the presence of any odor. Pups were then removed so that pouches with maternal and unfamiliar dam’s odors (10 g of bedding) could be added in the cylinders. They were then placed in the box again for a 10-minute testing period. The interactors and chambers were thoroughly cleaned with Virkon S solution in between trials.

### Play behavior

Play was assessed at four different points during the juvenile period to examine how behavior progressed as pups aged. One hour before play behavior was tested the pups were separated from each other in new cages. After the separation, pups were reunited in a dark room to promote play behavior in a clean test cage of dimensions 19 × 10 in^2^. Pups were placed in the center of the cage with their cage-mate and behavior was video-recorded for 10 minutes.

### Light-dark box test (LD)

For the LD test, rats were placed first in the light compartment of a two chambered custom-made apparatus (custom made, Curbell Plastics, Orchard Park, NY, USA) The juvenile apparatus (Plexiglas) was 16 × 40 × 21 cm^3^ and the adult apparatus (acrylic) was 100 × 100 × 40 cm^3^. The animals were allowed to freely explore both compartments for 5-minutes as previously described (Costall et al., 1989; Arrant et al., 2013). The time spent (Light Time, s) and distance travelled (Light, m) in the light compartment, the frequency of entering the light compartment (Light Entries, #), and the latency to enter the dark (Lat to Dark, s) compartment were analyzed using video-recordings in Ethovision. The apparatus was cleaned with Virkon S between trials.

### Social interaction test (SIT)

The SIT (Kaidanovich-Beilin et al., 2011) was performed in a custom-made 3-chambered social interaction apparatus (custom made, Curbell Plastics, Orchard Park, NY, USA; juvenile apparatus: white acrylic 42 × 62 × 30 cm^3^; adult apparatus: black acrylic 80 × 120 × 40 cm^3^). Rats were tested to determine their preference for a novel rat versus a novel object. Each was placed inside a cylinder with wired grid/mesh (juvenile interactors: 10 cm outer diameter, 25 cm height; adult interactors: 25 cm outer diameter, 25 cm height). Rats were first placed inside the center chamber and left to freely explore the entire apparatus with empty interactors located in the peripheral compartments for a 5-minute habituation period. Animals were then temporarily removed from the apparatus so that a novel rat and object could be placed inside the interactors. Placement of the novel rat or novel object were alternated between trials. Rats were then placed again in the central box of the apparatus and tested for 10-minutes. The box was cleaned with Virkon S between trials.

### Forced swim test (FST)

Adult animals (~P150) were tested in a two-day FST paradigm as described elsewhere (Slattery and Cryan, 2012; Yankelevitch-Yahav et al., 2015). The experiment consisted of an initial 15-minute training (pre-test) session on day 1, and a 5-minute testing session 24 hours later. A clear, Plexiglas container (custom made, Curbell Plastics, Orchard Park, NY, USA) with an internal diameter of 20 cm and a height of 50 cm was filled with tap water at a temperature of 24 ± 1°C, to a height such that the animal was unable to touch the bottom with its hind paws (35 ± 1 cm). After each trial, animals were dried with a towel by the experimenter and then placed in a clean holding cage atop a heating pad set to 37°C. Animals were reintroduced to their home cage when completely dry.

### Blood collection during FST for corticosterone

Each animal underwent blood draws at five distinct time points: immediately prior to the FST (time 0), and at 15 minutes, 30 minutes, 60 minutes, and 120 minutes post-FST. Blood was obtained via tail clipping and 20 to 100 μL at each time point was collected into a capillary tube with heparin (Microvette^®^ 100 K3E, EDTA Preparation, Sarstedt). The tube was immediately placed on ice. Serial samples of blood were collected at post-FST time points by manual disruption of the clot at the end of the tail. Pressure was applied to the wound for hemostasis. When blood flow was heavier than anticipated, styptic powder was applied. Following completion of the behavioral paradigm, the tubes were centrifuged at 5000 RPM for 10 minutes, in a 4°C cold room. The supernatant was then separated from the pellet with a pipette, and stored at −20°C for subsequent steroid analysis.

### Elevated plus maze (EPM)

Dams were tested in a standard plus-shaped EPM apparatus (custom made, Curbell Plastics, Orchard Park, NY, USA), consisting of two open arms and two closed arms of the same size (50 × 10 cm^2^) joined by a closed center square. The closed arms were enclosed by walls (35 cm) except at the joined platform. The entire apparatus was elevated to a height of 55 cm above the floor. Rats were placed individually at the center of the open field facing one of the open arms and were allowed to explore the apparatus for 10 minutes. The time spent in the open arms, number of entries into the open arms, and distance were measured. The EPM was cleaned with Virkon S between trials.

### Behavioral analysis

Behavioral data were collected using a high-definition webcam (C920, Logitech, Apples, Switzerland) or an infrared camera (EO-2223 NIR USB 3.0 Camera, Edmund Optics, Barrington, NJ, USA, 08007) when the behavior was conducted under infrared illumination (AT-11E-S 80, Axton Tech, North Salt Lake, UT 84054). Play behavior including durations (seconds) of rough-and-tumble (rolling around) was coded manually by an observer blind to the treatment groups using Avidemux software (version 2.7.4, http://www.avidemux.org) and Microsoft Excel. All other behaviors were analyzed with Ethovision XT software (version 15; Noldus Information Technology, Wageningen, the Netherlands), with the exception of climbing behavior in the FST, which was manually scored by an observer blind to the group and sex of the rat.

### Statistical Analysis

Data manipulation and graphing were performed in R using the `stringr`, `outliers`, `ggplot2`, and `dplyr` libraries while statistical analyses were done using the `rstatix` package. A 2 × 2 × 2 ANOVA was used to analyze the effects of Rearing, Sex and Housing or Rearing, Sex, and Testing. A repeated measures ANOVA was used to analyze time course data. Planned post-hoc comparisons were conducted using Tukey’s Test or Bonferroni’s correction to alpha for multiple, repeated measures tests. A two-tailed t-test was used to analyze EPM data. For corticosterone analyses, obtained absorbance data were analyzed in GraphPad Prism 8.0, where the serum corticosterone concentrations (ng/ml) were log-transformed and a two-way analysis of variance (ANOVA) with Bonferroni’s post-hoc test applied.

## Results

### A severe form of MS designed for maximal effects on biobehavioral profiles across the lifespan

Six-hours of maternal separation began on P2 and lasted until P21 (Fig. 1A). A total of 104 (C=52, MS=52, 1:1 male:female) rats were used in the study. Of these rats, 40 [C=20, MS=20, 1:1 male:female]) was randomly assigned to a cross-housed condition (control with maternally separated offspring). The remaining 64 rats were housed with siblings from the same litter (C=32, MS=32, 1:1 male:female). Of this group, 24 rats (C=12, MS=12,1:1 male:female) were randomly assigned to a non-tested group, which would not undergo juvenile testing. There were a total of 12 groups of rats across rearing, sex, housing, and testing conditions. Pup body weight was assessed at weaning (P21) and consistent with our previous results (Kaidbey et al., 2019), no significant differences in body weight between C and MS groups for either males or females was identified despite the prolonged separation (Fig. 1B). Interestingly, while there was also no effect of MS on the weight in adults (~P90), a rearing by sex interaction was identified with MS males weighting slightly less than their C counterparts. Furthermore, we assessed dam anxiety on the EPM test and found no differences on between MS and C dams (Fig. 1C).

### MS minimally affects juvenile behaviors and when it does, it is associated with protection against maladaptive behaviors

We first asked whether this severe form of MS reduces the pups’ preference for their own dam’s odor. The maternal odor preference (OP) test was conducted 1-4 days post-weaning (P22 to P25). We selected this time interval to allow the dam odor to accumulate in the newly changed cage after weaning and to impregnate the bedding to be used for the test. A 3-chambered arena was used with one side-chamber containing the juvenile’s own dam odor and the other side-chamber containing an unfamiliar dam odor. During the first 2-min all pus investigated the two odor-containing chambers equally. Therefore, we considered this to be a habituation period and focused analyses on the remaining 8 minutes of the test (Fig. 2A, left).

**Fig. 2.**
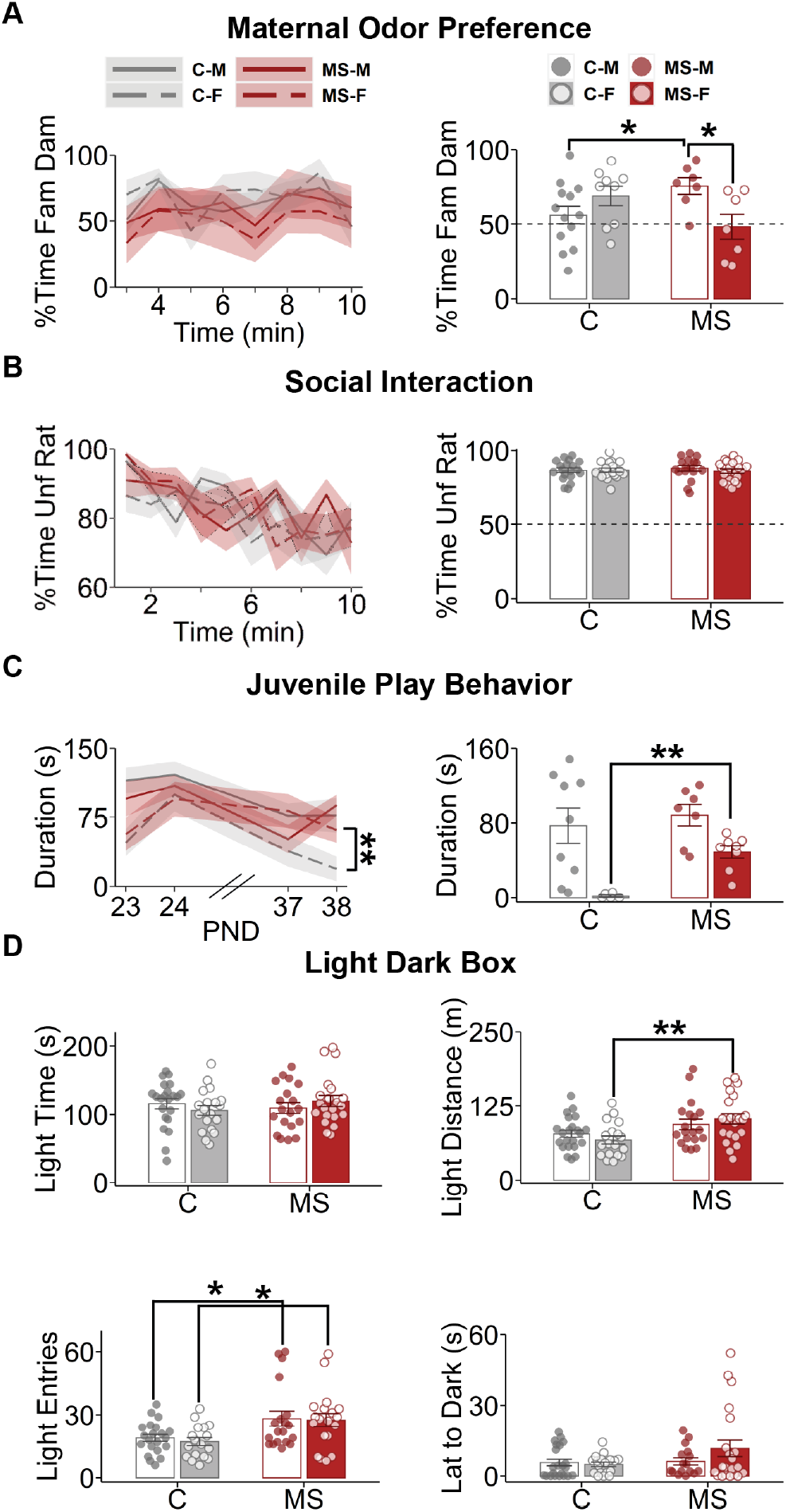
Effect of MS on maternal odor preference, juvenile sociability, and anxiety-like behavior. **(A)** Maternal odor preference was tested by presentation of the pup’s own dam’s odor versus an unfamiliar dam odor in a three-chamber arena. Data was analyzed for the last 8 min of 10 min of exploration. No significant differences were observed in 1-min binned time-series (left:% Time Fam Dam, Rearing: *F*_1,25_ = 1.67, *p* = 0.21, Sex: *F*_1,25_ = 0.16, *p* = 0.69, Time : *F*_7,175_ = 0.91, *p* = 0.50, Rearing × Sex: *F*_1,25_ = 0.246, *p* = 0.624, Rearing × Time: *F*_7,175_ = 1.36, *p* = 0.23, Sex × Time: *F*_7,175_ = 0.29, *p* = 0.96, Rearing × Sex × Time: *F*_7,175_ = 1.03, *p* = 0.41), but cumulatively, MS males did spent more time with their own dam’s odor than C males (right: % Time Fam Dam, Rearing: *F*_1,32_ = 0.008, *p* = 0.93, Sex: *F*_1,32_ = 1.09, *p* = 0.31, Rearing × Sex: *F*_1,32_ = 8.38, *p* = 0.007). **(B)**) No significant group differences were seen in the SIT over time (left: % Time Unf Rat, Rearing: *F*_1,67_ = 0.70, *p* = 0.40, Sex: *F*_1,67_ = 0.99, *p* = 0.32, Time: *F*_9,603_ = 4.93, *p* = 2.1 × 10^-6^, Rearing × Sex: *F*_1,67_ = 0.44, *p* = 0.51, Rearing × Time: *F*_9,603_ = 1.21, *p* = 0.29, Sex × Time: *F*_9,603_ = 0.99, *p* = 0.45, Rearing × Sex × Time : *F*_9,603_ = 1.17, *p* = 0.31) or cumulatively (right: % Time Unf Rat, Rearing: *F*_1,71_ = 0.036, *p* = 0.85, Sex: *F*_1,71_ = 0.34, *p* = 0.56, Rearing × Sex: *F*_1,71_ = 0.41, *p* = 0.53). **C** Repeated-measures ANOVA on the duration of time spent engaging in play behavior revealed an interaction between Rearing and Sex, which post-hoc testing showed to be driven by more play behavior in the MS females compared to C females on the last day of testing (left: Duration (s), Rearing: *F*_1,25_ = 1.58, *p* = 0.22, Sex: *F*_1,25_ = 26, *p* = 2.9 × 10^-5^, Time: *F*_3,75_ = 7.24, *p* = 2.5 × 10^-4^, Rearing × Sex: *F*_1,25_ = 8.42, *p* = 0.008, Rearing × Time: *F*_3,75_ = 0.77, *p* = 0.51, Sex × Time: *F*_3,75_ = 1.73, *p* = 0.17, Rearing × Sex × Time: *F*_3,75_ = 0.56, *p* = 0.64). An analysis of the last day alone confirmed this effect (right: Duration (s), Rearing: *F*_1,25_ = 4.36, *p* = 0.047, Sex: *F*_1,25_ = 17.1, *p* = 3.5 × 10^-4^, Rearing × Sex: *F*_1,25_ = 1.65, *p* = 0.21). **D**Light dark box testing revealed less anxiety-like behavior in MS rats on several measures: light time (top left: Light Time (s), Rearing: *F*_1,77_ = 0.24, *p* = 0.63, Sex: *F*_1,77_ = 7.0 × 10^-4^, *p* = 0.98, Rearing × Sex: *F*_1,77_ = 1.70, *p* = 0.19), distance travelled in the light zone (top right: Light Distance (m), Rearing: *F*_1,76_ = 11.3, *p* = 0.001, Sex: *F*_1,76_ = 0.007, *p* = 0.94, Rearing × Sex: *F*_1,76_ = 1.71, *p* = 0.20), light transitions or entries (lower left: Light Entries, Rearing: *F*_1,76_ = 11.3, *p* = 0.001, Sex: *F*_1,76_ = 0.007, *p* = 0.94, Rearing × Sex: *F*_1,76_ = 1.71, *p* = 0.20), latency to enter the dark zone (lower right: Lat to Dark (s), Rearing: *F*_1,73_ = 2.65, *p* = 0.11, Sex: *F*_1,73_ = 1.13, *p* = 0.29, Rearing × Sex: *F*_1,73_ = 1.99, *p* = 0.16). Fam, familiar; Unf, unfamiliar; Lat, latency; Light, light zone; Dark, dark zone; MS, maternally separated; C, control. Bars represent means ± SEM. Lines represent means. Shaded regions represent SEM. (Tukey’s or Bonferroni’s tests, \**p* < 0.05, \*\**p* < 0.01).

We found no significant differences between groups on repeated measures ANOVA on 1-min binned exploration (Rearing: *F*_1,25_ = 1.67, *p* = 0.21, Sex: *F*_1,25_ = 0.16, *p* = 0.69, Time: *F*_7,175_ = 0.91, *p* = 0.50, Rearing × Sex: *F*_1,25_ = 0.25, *p* = 0.62, Rearing × Time: *F*_7,175_ = 1.36, *p* = 0.23, Sex × Time: *F*_7,175_ = 0.29, *p* = 0.96, Rearing × Sex × Time: *F*_7,175_ = 1.03, *p* = 0.41). However, a 2 × 2 ANOVA on cumulative exploration during the 8min revealed a significant interaction between Rearing and Sex (Rearing: *F*_1,32_ = 0.008, *p* = 0.93, Sex: *F*_1,32_ = 1.09, *p* = 0.31, Rearing × Sex: *F*_1,32_ = 8.38, *p* = 0.007) (Fig. 2A, right), with MS male but not female pups spending significantly more time investigating their own dam’s odors compared to both C males (Tukey’s *p* = 0.049) and MS female (Tukey’s *p* = 0.019).

Next, we evaluated juvenile social behaviors. Social interaction and juvenile play were tested between P25 and P45. We first used the three-chambered SIT to quantify social preference toward an unfamiliar rat versus a novel object. No significant differences were found between C and MS (Rearing: *F*_1,67_ = 0.70, *p* = 0.40, Sex: *F*_1,67_ = 0.99, *p* = 0.32, Time: *F*_9,603_ = 4.93, *p* < 1 × 10^-5^, Rearing × Sex: *F*_1,67_ = 0.44, *p* = 0.51, Rearing × Time: *F*_9,603_ = 1.21, *p* = 0.29, Sex × Time: *F*_9,603_ = 0.99, *p* = 0.45, Rearing × Sex × Time: *F*_9,603_ = 1.17, *p* = 0.31) (Fig. 2B, left). Both C and MS groups preferred the unfamiliar rat, spending ~85% of their time in social exploration (Rearing: *F*_1,71_ = 0.036, *p* = 0.85, Sex: *F*_1,71_ = 0.34, *p* = 0.56, Rearing × Sex: *F*_1,71_ = 0.41, *p* = 0.53) (Fig 2B, right). Juvenile play was tested on 2 consecutive days twice, at P23-P24 and P37-P38.

A repeated-measures ANOVA on the duration of time spent engaging in play behavior revealed an interaction between Rearing and Sex (Rearing: *F*_1,25_ = 1.58, *p* = 0.22, Sex: *F*_1,25_ = 26, *p* = 2.9 × 10^-5^, Time: *F*_3,75_ = 7.24, *p* = 2.5 × 10^-4^, Rearing × Sex: *F*_1,25_ = 8.42, *p* = 0.008, Rearing × Time: *F*_3,75_ = 0.77, *p* = 0.51, Sex × Time: *F*_3,75_ = 1.73, *p* = 0.17, Rearing × Sex × Time: *F*_3,75_ = 0.56, *p* = 0.64) (Fig. 2C, left). Bonferroni post-hoc test showed that the effects of Sex and its interaction with Rearing were due to more play behavior in the MS females compared to C females on the last day of testing (Bonferroni’s *p* = 0.0015). An analysis of the last day (Rearing: *F*_1,25_ = 4.36, *p* = 0.047, Sex: *F*_1,25_ = 17.1, *p* = 3.5 × 10^-4^, Rearing × Sex: *F*_1,25_ = 1.65, *p* = 0.21) alone confirmed this effect (Tukey’s *p* =1.9 × 10^-4^) (Fig 2C, right).

We then assessed the effect of MS on anxiety-like behavior in the LD test (Fig. 2D). No significant differences between groups were observed in time spent in the light zone (Rearing: *F*_1,77_ = 0.24, *p* = 0.63, Sex: *F*_1,77_ = 7 × 10^-4^, *p* = 0.98, Rearing × Sex: *F*_1,77_ = 1.70, *p* = 0.19). However, there was a significant effect of Rearing was observed in the distance traveled in the light zone, with MS rats traveling a greater distance than C rats (Rearing: *F*_1,76_ = 11.3, *p* = 0.001, Sex: *F*_1,76_ = 0.007, *p* = 0.94, Rearing × Sex: *F*_1,76_ = 1.71, *p* =0.20), and this this effect was largely driven by female rats (Tukey’s *p* = 0.0039). Additionally, both male and female MS rats also had significantly more entries into the light zone (Rearing: *F*_1,76_ = 11.3, *p* = 0.001, Sex: *F*_1,76_ = 0.007, *p* = 0.94, Rearing × Sex: *F*_1,76_ = 1.71, *p* = 0.20, males Tukey’s *p* = 0.019, females Tukey’s *p* = 0.010), overall suggesting MS is associated with a lower anxiety-like phenotype.

### MS effects on sociability and anxiety-like behavior in the adult period mostly mirror juvenile phenotypes

MS effects on sociability and anxiety-like behavior in the adult period mostly mirror juvenile phenotypes Social preference was examined again using SIT between P90 and P150 to evaluate for possible emergence of changes in sociability. Strong social preference was again seen across all groups. No significant effect of Rearing was observed (Rearing: *F*_1,98_ = 0.004, *p* = 0.95, Sex: *F*_1,98_ = 18.9, *p* = 3.3 × 10^-5^, Rearing × Sex: *F*_1,98_ = 0.014, *p* = 0.91), although a significant effect of Sex was identified, with males spending more time with an unfamiliar rat than females (80% vs 72%, Tukey’s *p* = 2.7 × 10^-5^) (Fig. 3A, right). Time series analysis using repeated measures ANOVA revealed significant effects of Sex and Time (Rearing: *F*_1,91_ = 0.011, *p* = 0.92, Sex: *F*_1,91_ =21, *p* = 1.4 × 10^-5^, Time: *F*_9,819_ = 4.31, *p* =1.7 × 10^-5^, Rearing × Sex: *F*_1,91_ = 0.012, *p* = 0.92, Rearing × Time: *F*_9,819_ = 1.82, *p* = 0.62, Sex × Time: *F*_9,819_ = 1.28, *p* = 0.24, Rearing × Sex × Time: *F*_9,819_ = 0.73, *p* = 0.68). Bonferroni post-hoc tests did not show significant differences between groups at any particular time but did show that the preference for the unfamiliar rat was significantly higher in the first minute compared to the rest of the 9 minutes of the experiment (Fig. 3A, left).

**Fig. 3.**
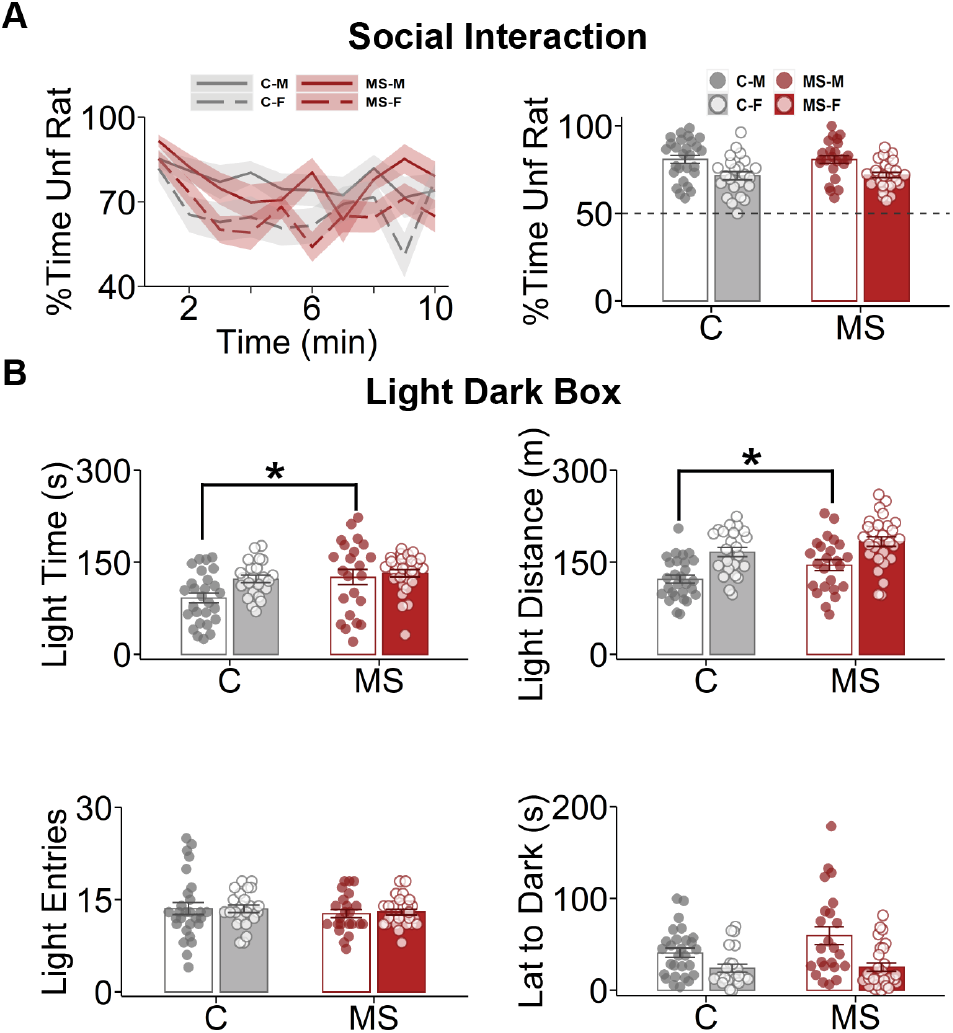
Effect of MS on adult sociability and anxiety-like behavior mostly mirror juvenile phenotypes. **(A)** No effects of MS were observed on SIT though significant Sex effects were observed both in time series analysis (left: % Time Unf Rat, Rearing: *F*_1,91_ = 0.011, *p* = 0.92, Sex: *F*_1,91_ =21, *p* = 1.8 × 10^-5^, Time: *F*_9,819_ = 4.31, *p* = 1.7 × 10^-5^, Rearing × Sex: *F*_1,91_ = 0.012, *p* = 0.92, Rearing × Time: *F*_9,819_ = 1.82, *p* = 0.62, Sex × Time: *F*_9,819_ = 1.28, *p* = 0.24, Rearing × Sex × Time: *F*_9,819_ = 0.73, *p* = 0.68) and cumulative time spent with an unfamiliar rat (right: % Time Unf Rat, Rearing: = 0.004, *p* = 0.95, Sex: *F*_1,98_ = 18.9, *p* = 3.3 × 10^-5^, Rearing × Sex: *F*_1,98_ = 0.014, *p* = 0.91), with males spending more time in social exploration than females (Tukey’s *p* = 2.7 × 10^-5^). **(B)** Light dark box testing revealed significant effects of Rearing, with MS males spending more time and traveling a longer distance in the light zone than C males, and Sex, with females having shorter latency to the dark zone, but spending more time in, traveling longer distance in, and showing more transitions to the light zone. (top left: Light Time (s), Rearing: *F*_1,97_ = 6.90, *p* = 0.01, Sex: *F*_1,97_ = 5.14, *p* = 0.026, Rearing × Sex: *F*_1,97_ = 2.26, *p* = 0.14; top right: Light Distance (m), Rearing: *F*_1,99_ = 6.76, *p* = 0.011, Sex: *F*_1,99_ = 29, *p* = 5.1 × 10^-7^, Rearing × Sex: *F*_1,99_ = 0.15, *p* = 0.70; lower left: Light Entries, Rearing: *F*_1,97_ = 0.86, *p* = 0.36, Sex: *F*_1,97_ = 0.021, *p* = 0.89, Rearing × Sex: *F*_1,97_ = 0.045, *p* = 0.83; lower right: Lat to Dark (s), Rearing: *F*_1,97_ = 2.70, *p* = 0.10, Sex: *F*_1,97_ = 17.7, *p* = 5.7 × 10^-5^, Rearing × Sex: *F*_1,97_ = 2.05, *p* = 0.16). Fam, familiar; Unf, unfamiliar; Lat, latency; Light, light zone; Dark, dark zone. Bars represent means ± SEM. Lines represent means. Shaded regions represent SEM. (Tukey’s or Bonferroni’s tests, \**p* < 0.05, \*\**p* < 0.01).

Anxiety-like behavior was reexamined using LD testing. A similar effect of Rearing (Rearing: *F*_1,97_ = 6.90, *p* = 0.01, Sex: *F*_1,97_ = 5.14, *p* = 0.026, Rearing × Sex: *F*_1,97_ = 2.26, *p* = 0.14) was observed as during the juvenile period, with MS males spending more time (Tukey’s *p* = 0.021) and traveling a longer distance (Rearing: *F*_1,99_ = 6.76, *p* = 0.011, Sex: *F*_1,99_ = 28.87, *p* = 5.1 × 10^-7^, Rearing × Sex: *F*_1,99_ = 0.15, *p* = 0.70) in the light zone than C males (Tukey’s *p* = 0.037) (Fig. 3B). A significant effect of Sex was also observed (Rearing: *F*_1,97_ = 2.70, *p* = 0.10, Sex: *F*_1,97_ = 17.7, *p* = 5.8 × 10^-5^, Rearing × Sex: *F*_1,97_ = 2.05, *p* = 0.16), with females having shorter latency to the dark zone (Tukey’s *p* = 0.0099) but spending more time in (Tukey’s *p* = 0.018), and traveling longer distance in (Tukey’s *p* < 3.1 × 10^-7^) the light zone. However, no significant group differences were found in transitions to the light zone (Rearing: *F*_1,97_ = 0.86, *p* = 0.36, Sex: *F*_1,97_ = 0.021, *p* = 0.89, Rearing × Sex: *F*_1,97_ = 0.045, *p* = 0.83).

### MS is associated with long term negative coping-behavioral outcomes but not differences in blood corticosterone responses

The FST (Porsolt test) was used to evaluate effects of MS on coping behaviors. A 15-minute pretest period the day before the testing period (5-min) was used to habituate the rats (Fig. 4A). Additionally, baseline corticosterone was collected before testing began on Day 2 (T0), as well as 15 minutes (T15), 30 minutes (T30), 60 minutes (T60), and 120 minutes (T120) after the FST. The FST measures that were analyzed were: immobility time, latency to the first immobility state, climbing time, and struggling time.

**Fig. 4.**
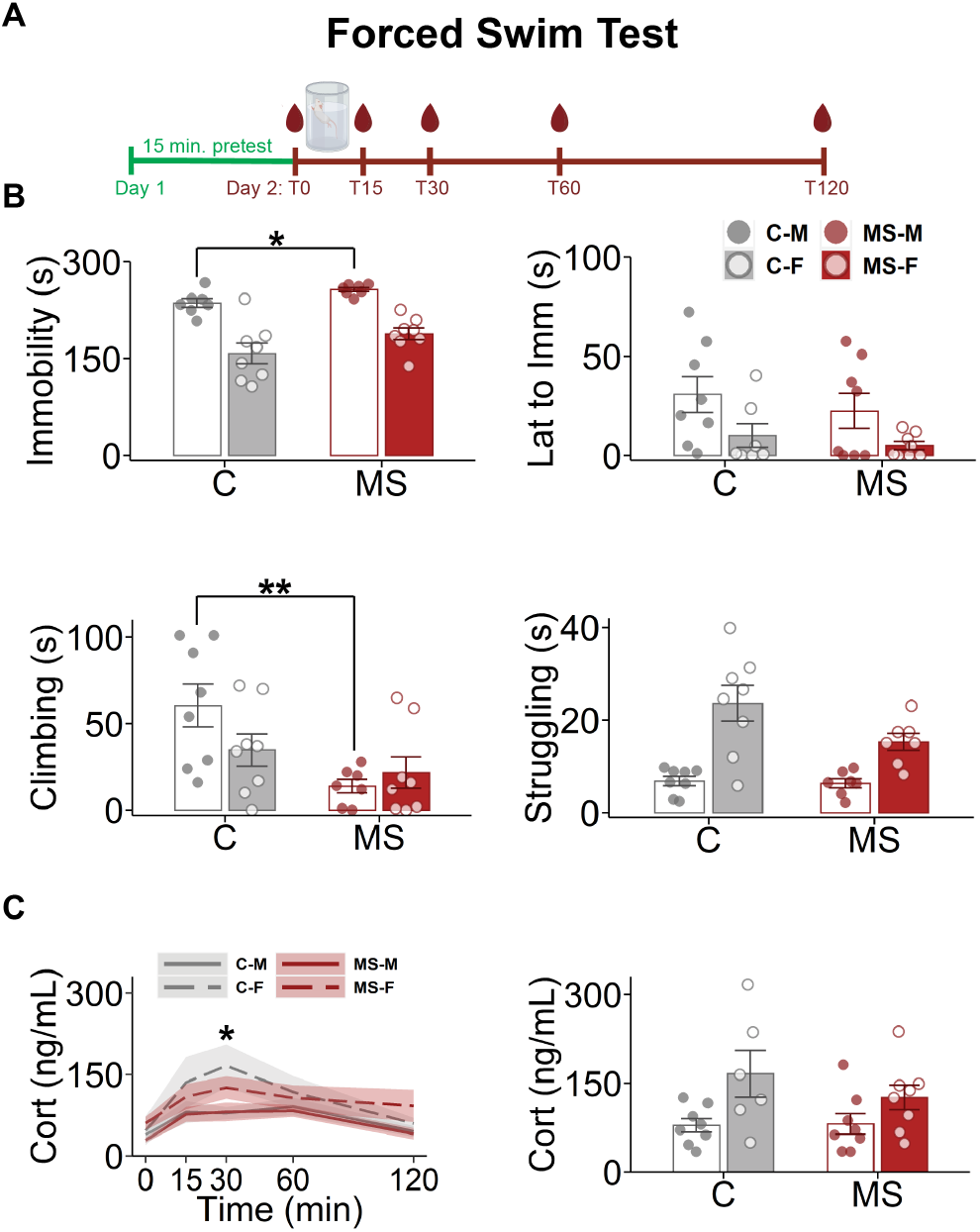
Effect of MS on adult sociability and anxiety-like behavior mostly mirror juvenile phenotypes. **(A)** Experimental timeline for the forced swim and blood corticosterone collection timeline. A 15-minute pretest was done on the day before the testing began (5-minute test). Blood was collected at baseline (T0), 15 minutes (T15), 30 minutes (T30), 60 minutes (T60) and 120 minutes (T120) after baseline. **(B)** Rearing and Sex effects were identified on several measures of the FST (top left: Immobility (s), Rearing: *F*_1,26_ = 5.98, *p* = 0.022, Sex: *F*_1,26_ = 49, *p* = 1.8 × 10^-7^, Rearing × Sex: *F*_1,26_ = 0.21, *p* = 0.65; top right: Lat to Imm (s), Rearing: *F*_1,27_ = 0.89, *p* = 0.36, Sex: *F*_1,27_ = 7.16, *p* = 0.012, Rearing × Sex: *F*_1,27_ = 0.82, *p* = 0.82; middle left: Climbing (s), Rearing: *F*_1,27_ = 9.89, *p* = 0.004, Sex: *F*_1,27_ = 0.89, *p* = 0.36, Rearing × Sex: *F*_1,27_ = 3.15, *p* = 0.08; middle right: Struggling (s), Rearing: *F*_1,26_ = 3.57, *p* = 0.70, Sex: *F*_1,26_ = 30, *p* = 9.6 × 10^-6^, Rearing × Sex: *F*_1,26_ = 2.80, *p* = 0.11), with MS males spending more time immobile (Tukey’s *p* = 0.018), and females exhibiting shorter latencies to immobility (Tukey’s *p* = 0.010) but spending overall less time immobile (Tukey’s *p* = 4.2 × 10^-7^) and more time struggling (Tukey’s *p* = 1.6 × 10^-5^) compared to males. **(C)** Serum corticosterone levels revealed the expected increase immediately after FST but did not reveal a significant Rearing or Sex effects (Cort [ng/mL], Rearing: *F*_1,19_ = 0.15, *p* = 0.70 Sex: *F*_1,19_ = 1.93, *p* = 0.18, Time: *F*_9,819_ = 4.31, *p* = 6.4 × 10^-10^, Rearing × Sex: *F*_1,91_ = 0.012, *p* = 0.92, Rearing × Time: *F*_9,819_ = 1.82, *p* = 0.62, Sex × Time : *F*_9,819_ = 1.28, *p* = 0.24, Rearing × Sex × Time: *F*_9,819_ = 0.73, *p* = 0.68). Post hoc testing revealed corticosterone was significantly higher than baseline at T15, T30 and T60, but not T120 (Bonferroni, T15 *p* = 0.007; T30 *p* = 2.9 × 10^-4^; T60 *p* = 3.8 × 10^-4^). Additionally, post hoc testing revealed a Sex effect at T30, with females having a significantly higher corticosterone level than males (Bonferroni, adjusted *p* = 0.04). This Sex effect was confirmed on 2×2 ANOVA on the corticosterone levels at T30 (Rearing: *F*_1,26_ = 0.71, *p* = 0.41, Sex: *F*_1,26_ = 8.81, *p* = 0.006, Rearing × Sex: *F*_1,26_ = 0.91, *p* = 0.35). Bars represent means ± SEM. Lines represent means. Shaded regions represent SEM. (Tukey’s, \**p* < 0.05, \*\**p* < 0.01).

A 2 × 2 ANOVA revealed significant effects of Rearing and Sex on immobility (*F*_1,26_ = 5.98, *p* = 0.022, Sex: *F*_1,26_ = 49, *p* = 1.8 × 10^-7^, Rearing × Sex: *F*_1,26_ = 0.21, *p* = 0.65) (Fig. 4B). Post-hoc analysis revealed this effect was largely driven by MS males, which spent significantly more time immobile (257 ± 3 s) than C males (236 ± 7 s) (Tukey’s *p* = 0.018) and females (C: 158 ± 16 s, MS: 188 ± 9 s) (Fig. 4B). An effect of Sex but not Rearing was observed on the latency to the first immobility state (Rearing: *F*_1,27_ = 0.89, *p* = 0.36, Sex: *F*_1,27_ = 7.16, *p* = 0.012, Rearing × Sex: *F*_1,27_ = 0.82, *p* = 0.82), with females becoming immobile faster (Tukey’s *p* = 0.010).

Climbing behaviors, which can be indicative of a positive coping behavioral strategy, showed significant main effect of Rearing (Rearing: *F*_1,27_ = 9.89, *p* = 0.004, Sex: *F*_1,27_ = 0.89, *p* = 0.36, Rearing × Sex: *F*_1,27_ = 3.15, *p* = 0.08). Mirroring the results observed in immobility, this effect was driven mainly by MS males spending less time climbing compared with C males (13.9 ± 4.0 s versus 60.5 ± 12.4 s, respectively), as confirmed by a planned post-hoc testing (Tukey’s *p* = 0.0050). Struggling only revealed a significant effect of Sex (Rearing: *F*_1,26_ = 3.57, *p* = 0.70, Sex: *F*_1,26_ = 30, *p* = 9.6 × 10^-6^, Rearing × Sex: *F*_1,26_ = 2.80, *p* = 0.11), with female rats spending more time struggling than male rats (Tukey’s *p* = 1.6 × 10^-5^).

Above-described MS effects on FST are unlikely to be mediated by differences in stress-hormones, as no significant differences were identified on blood corticosterone levels except for Time (Rearing: *F*_1,19_ = 0.15, *p* = 0.70 Sex: *F*_1,19_ = 1.93, *p* = 0.18, Time : *F*_9,819_ = 4.31, *p* = 6.4 × 10^-10^, Rearing × Sex: *F*_1,91_ = 0.012, *p* = 0.92, Rearing × Time : *F*_9,819_ = 1.82, *p* = 0.62, Sex × Time : *F*_9,819_ = 1.28, *p* = 0.24, Rearing × Sex × Time : *F*_9,819_ = 0.73, *p* = 0.68), which revealed corticosterone concentrations to increase during the FST compared to baseline (T0), except at T120 (Bonferroni, adjusted p, T15, *p* = 0.007; T30 *p* = 2.9 × 10^-4^; T60 *p* = 3.8 × 10^-4^) and to decrease between T30, T60 and T120 (Bonferroni, adjusted *p*, T60, *p* = 0.016; T120 *p* = 0.042).

Post hoc analyses also revealed a significant Sex effect at 30 min, with females having significantly higher corticosterone levels compared to males (Bonferroni, adjusted *p* = 0. 04). This was confirmed by a 2 × 2 ANOVA on the corticosterone levels at this timepoint (Rearing: *F*_1,26_ = 0.71, *p* = 0.41, Sex: *F*_1,26_ = 8.81, *p* = 0.006, Rearing × Sex: *F*_1,26_ = 0.91, *p* = 0.35).

### Minimal effects observed as a function of post-weaning housing with sibling versus unfamiliar rat

Post-weaning housing conditions can play a pivotal role in neurodevelopment (Smith et al., 1997) and potentially interact with preweaning MS. We therefore investigated whether adult behavioral outcomes were influenced by post-weaning housing with sibling versus unfamiliar rat from a cross-housing condition (control with maternally separated). No effects of Housing were observed on the SIT (Rearing: *F*_1,94_ = 0.036, *p* = 0.85, Sex: *F*_1,94_ = 21, *p* = 1.6 × 10^-5^, Housing: *F*_1,94_ = 0.66, *p* = 0.42, Rearing × Sex: *F*_1,94_ = 0.027, *p* = 0.87, Rearing × Housing: *F*_1,94_ = 0.59, *p* = 0.44, Sex × Housing: *F*_1,94_ = 2.29, *p* = 0.13, Rearing × Sex × Housing: *F*_1,94_ = 0.067, *p* = 0.79) (Fig 5A). Housing effects were also not observed in LD testing on time spent in (Rearing: *F*_1,93_ = 6.53, *p* = 0.012, Sex: *F*_1,93_ = 4.16, *p* = 0.044, Housing: *F*_1,93_ = 0.45, *p* = 0.50, Rearing × Sex: *F*_1,93_ = 1.54, *p* = 0.22, Rearing × Housing: *F*_1,93_ = 0.066, *p* = 0.79, Sex × Housing: *F*_1,93_ = 0.51, *p* = 0.48, Rearing × Sex × Housing: *F*_1,93_ = 0.47, *p* = 0.49), distance traveled in (Rearing: *F*_1,95_ = 6.51, *p* = 0.012, Sex: *F*_1,95_ = 27, *p* = 1.2 × 10^-6^, Housing: *F*_1,95_ = 2.13, *p* = 0.15, Rearing × Sex: *F*_1,95_ = 0.066, *p* = 0.79, Rearing × Housing: *F*_1,95_ = 0.077, *p* = 0.78, Sex × Housing : *F*_1,95_ = 2.0 × 10^-4^, *p* = 0.99, Rearing × Sex × Housing : *F*_1,95_ = 0.030, *p* = 0.86) or entries into the light chamber (Rearing: *F*_1,93_ = 1.37, *p* = 0.25, Sex: *F*_1,93_ = 2.0 × 10^-4^, *p* = 0.99, Housing: *F*_1,93_ = 0.30, *p* = 0.59, Rearing × Sex: *F*_1,93_ = 0.002, *p* = 0.96, Rearing × Housing: *F*_1,93_ = 1.21, *p* = 0.28, Sex × Housing: *F*_1,93_ = 0.15, *p* = 0.70, Rearing × Sex × Housing: *F*_1,93_ = 0.54, *p* = 0.46). A significant effect of Housing was observed on latency to enter the dark zone of the LD test (Rearing: *F*_1,94_ = 2.64, *p* = 0.11, Sex: *F*_1,94_ = 14.2, *p* = 2.9 × 10^-4^, Housing: *F*_1,94_ = 7.58, *p* = 0.007, Rearing × Sex: *F*_1,94_ = 1.44, *p* = 0.23, Rearing × Housing: *F*_1,94_ = 3.33, *p* = 0.07, Sex × Housing: *F*_1,94_ = 2.28, *p* = 0.09, Rearing × Sex × Housing: *F*_1,94_ = 1.01, *p* = 0.32) (Fig. 5B). Post-hoc tests revealed that MS males who were sibling-housed had longer latency to enter the dark zone compared with sibling-housed C males (Tukey’s *p* = 0.023) and MS male crossed-housed rats (Tukey’s *p* = 0.020).

**Fig. 5.**
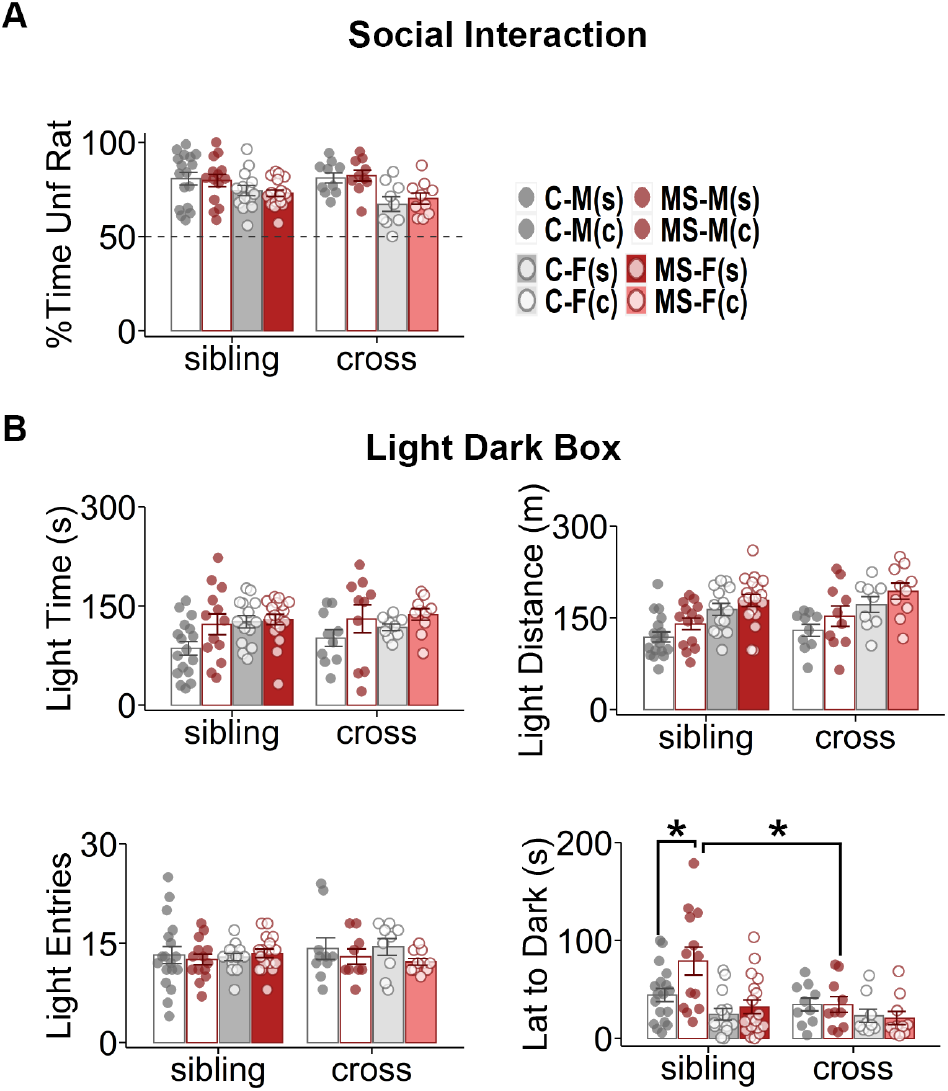
Minimal effects observed as a function of post-weaning housing with sibling versus unfamiliar rat. **(A)**Post-weaning housing with sibling versus an cross-housing condition (control with MS) had no effect on SIT (% Time Unf Rat: Rearing: *F*_1,94_ = 0.036, *p* = 0.85, Sex: *F*_1,94_ = 21, *p* = 1.6 × 10^-5^, Housing: *F*_1,94_ = 0.66, *p* = 0.42, Rearing × Sex: *F*_1,94_ = 0.027, *p* = 0.87, Rearing × Housing: *F*_1,94_ = 0.59, *p* = 0.44, Sex × Housing: *F*_1,94_ = 2.29, *p* = 0.13, Rearing × Sex × Housing: *F*_1,94_ = 0.067, *p* = 0.79). **(B)** Housing effects were also not observed in LD testing on time spent in, distance traveled in or entries into the light chamber, but a significant effect of Housing was observed on latency to enter the dark zone of the LD test (top left: Light Time (s): Rearing: *F*_1,93_ = 6.53, *p* = 0.012, Sex: *F*_1,93_ = 4.16, *p* = 0.044, Housing: *F*_1,93_ = 0.45, *p* = 0.50, Rearing × Sex: *F*_1,93_ = 1.54, *p* = 0.22, Rearing × Housing: *F*_1,93_ = 0.066, *p* = 0.79, Sex × Housing: *F*_1,93_ = 0.51, *p* = 0.48, Rearing × Sex × Housing: *F*_1,93_ = 0.47, *p* = 0.49; top right: Light Distance (m): Rearing: *F*_1,95_ = 6.51, *p* = 0.042, Sex: *F*_1,95_ = 27, *p* = 3.7 × 10^-5^, Housing: *F*_1,95_ = 2.13, *p* = 0.15, Rearing × Sex: *F*_1,95_ = 0.066, *p* = 0.79, Rearing × Housing: *F*_1,95_ = 0.077, *p* = 0.78, Sex × Housing: *F*_1,95_ = 2.0 × 10^-4^, *p* = 0.99, Rearing × Sex × Housing: *F*_1,95_ = 0.030, *p* = 0.86; lower left: Light Entries, Rearing: *F*_1,93_ = 1.37, *p* = 0.25, Sex: *F*_1,93_ = 2.0 × 10^-4^, *p* = 0.99, Housing: *F*_1,93_ = 0.30, *p* = 0.59, Rearing × Sex: *F*_1,93_ = 0.002, *p* = 0.96, Rearing × Housing: *F*_1,93_ = 1.21, *p* = 0.28, Sex × Housing: *F*_1,93_ = 0.15, *p* = 0.70, Rearing × Sex × Housing: *F*_1,93_ = 0.54, *p* = 0.46; Lat to Dark (s): Rearing: *F*_1,94_ = 2.64, *p* = 0.11, Sex: *F*_1,94_ = 14.2, *p* = 2.9 × 10^-4^, Housing: *F*_1,94_ = 7.58, *p* = 0.007, Rearing × Sex: *F*_1,94_ = 1.44, *p* = 0.23, Rearing × Housing: *F*_1,94_ = 3.33, *p* = 0.07, Sex × Housing: *F*_1,94_ = 2.28, *p* = 0.09, Rearing × Sex × Housing: *F*_1,94_ = 1.01, *p* = 0.32). Fam, familiar; Unf, unfamiliar; Lat, latency; Light, light zone; Dark, dark zone. Bars represent means ± SEM. Lines represent means. Shaded regions represent SEM. (Tukey’s, \**p* < 0.05, \*\**p* < 0.01).

### Testing during the juvenile period influences adult behavioral outcomes in MS rats

Test-retest reliability has been demonstrated to be low in rodent behavioral analysis (Andreatini and Bacellar, 2000) and significant habituation effects due to both repeat testing and cumulative handling have been observed (Bronstein et al., 1974; Gouveia and Hurst, 2017). Because not all rats were tested in the juvenile period, we therefore asked if juvenile testing influenced results of adult behavioral outcomes.

A significant effect of Sex and Prior Testing was observed in the SIT (Rearing: *F*_1,94_ = 0.27, *p* = 0.61, Sex: *F*_1,94_ = 16.8, *p* = 8.9 × 10^-5^, Testing: *F*_1,94_ = 4.37, *p* = 0.039, Rearing × Sex: *F*_1,94_ = 1.14, *p* = 0.29, Rearing × Testing: *F*_1,94_ = 1.37, *p* = 0.25, Sex × Testing: *F*_1,94_ = 0.46, *p* = 0.50, Rearing × Sex × Testing: *F*_1,94_ = 3.94, *p* = 0.05), with post-hoc analysis revealing untested C males exhibited a stronger social preference toward an unfamiliar rat than previously tested C males and untested MS males (Tukey’s *p* = 0.0044) (Fig. 6A). Prior Testing also was associated with more sociability in MS males compared to MS females (Tukey’s *p* = 0.0023).

**Fig. 6.**
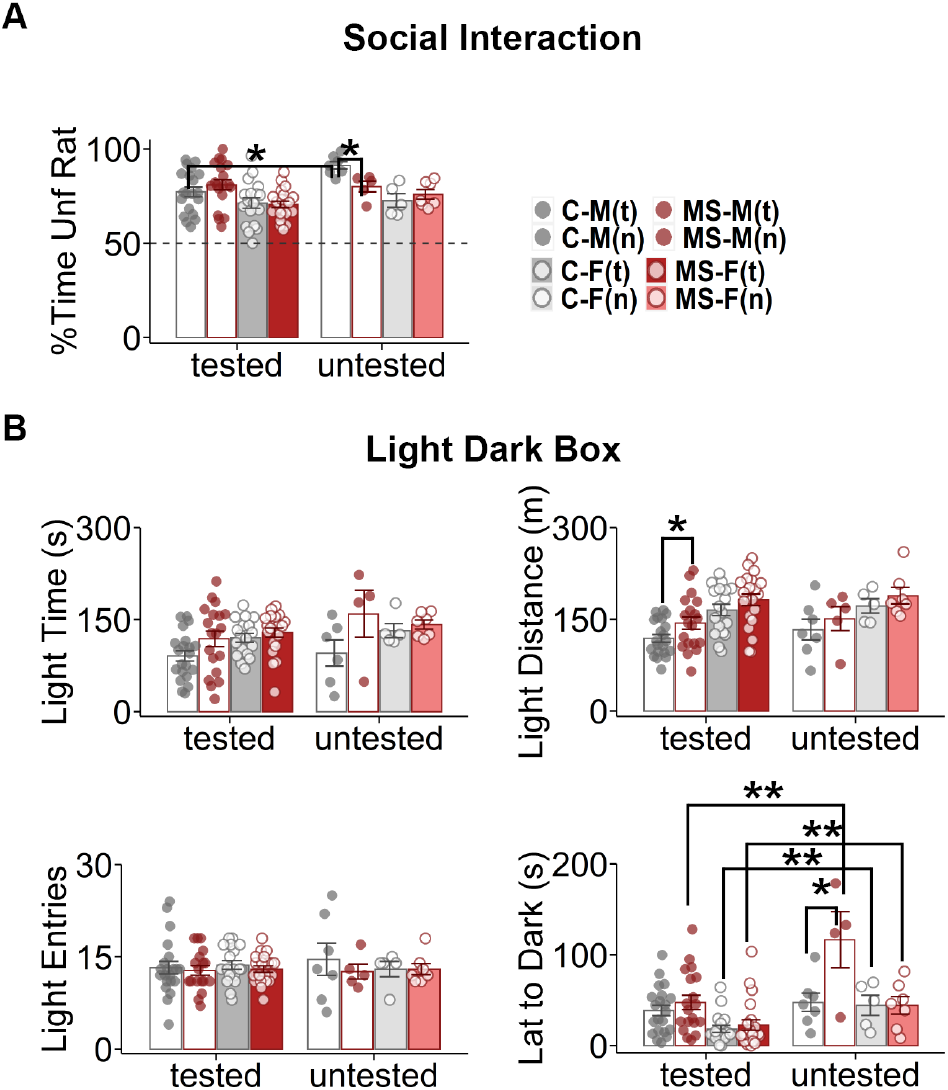
Testing during the juvenile period influences adult behavioral outcomes in MS rats. **(A)**A significant effect of Sex and Prior Testing was observed in the SIT, with post-hoc analysis revealing untested C males exhibited a stronger social preference toward an unfamiliar rat than previously tested C males and untested MS males. Prior Testing also was associated with more sociability in MS males compared to MS females (% Time Unf Rat: Rearing: *F*_1,94_ = 0.27, *p* = 0.61, Sex: *F*_1,94_ = 16.8, *p* = 8.8 × 10^-5^, Testing: *F*_1,94_ = 4.37, *p* = 0.039, Rearing × Sex: *F*_1,94_ = 1.14, *p* = 0.29, Rearing × Testing: *F*_1,94_ = 1.37, *p* = 0.25, Sex × Testing: *F*_1,94_ = 0.46, *p* = 0.50, Rearing × Sex × Testing: *F*_1,94_ = 3.94, *p* = 0.05). **(B)**LD testing was also influenced by prior testing, with significant effects observed by Rearing and Prior Testing, and interactions between Rearing and Sex, Rearing and Prior Testing, and Rearing, Sex, and Prior Testing on the latency to enter the dark zone in the LD test. Untested MS males took a significantly longer time to enter the dark zone than tested MS males. This was also true for untested female rats regardless of Rearing condition. (top left: Light Time (s): Rearing: *F*_1,93_ = 7.43, *p* = 0.008, Sex: *F*_1,93_ = 2.09, *p* = 0.15, Testing: *F*^1,93^ = 2.98, *p* = 0.08, Rearing × Sex: *F*_1,93_ = 3.30, *p* = 0.07, Rearing × Testing: *F*_1,93_ = 0.84, *p* = 0.36, Sex × Testing: *F*_1,93_ = 0.27, *p* = 0.60, Rearing × Sex × Testing: *F*_1,93_ = 0.73, *p* = 0.39; top right: Light Distance (m): Rearing: *F*_1,95_ = 4.23, *p* = 0.042, Sex: *F*_1,95_ = 18.8, *p* = 3.7 × 10^-5^, Testing: *F*_1,95_ = 0.88, *p* = 0.35, Rearing × Sex: *F*_1,95_ = 0.062, *p* = 0.80, Rearing × Testing: *F*_1,95_ = 0.038, *p* = 0.84, Sex × Testing : *F*_1,95_ = 0.045, *p* = 0.83, Rearing × Sex × Testing: *F*_1,95_ = 0.038, *p* = 0.85; lower left: Light Entries, Rearing: *F*_1,93_ = 0.78, *p* = 0.38, Sex: *F*_1,93_ = 0.022, *p* = 0.88, Testing: *F*_1,93_ = 0.019, *p* = 0.89, Rearing × Sex: *F*_1,93_ = 0.25, *p* = 0.62, Rearing × Testing: *F*_1,93_ = 0.058, *p* = 0.81, Sex × Testing: *F*_1,93_ = 0.27, *p* = 0.61, Rearing × Sex × Testing: *F*_1,93_ = 0.39, *p* = 0.54; Lat to Dark (s): Rearing: *F*_1,94_ = 9.06, *p* = 0.003, Sex: *F*_1,94_ = 19.5, *p = 2.7* × 10^-5^, Testing: *F*_1,94_ = 21, *p* = 1.4 × 10^-5^, Rearing × Sex: *F*_1,94_ = 7.26, *p* = 0.008, Rearing × Testing: *F*_1,94_ = 4.16, *p* = 0.044, Sex × Testing: *F*_1,94_ = 1.21, *p* = 0.28, Rearing × Sex × Testing: *F*_1,94_ = 5.54, *p* = 0.021). Fam, familiar; Unf, unfamiliar; Lat, latency; Light, light zone; Dark, dark zone. Bars represent means ± SEM. Lines represent means. Shaded regions represent SEM. (Tukey’s, \**p* < 0.05, \*\**p* < 0.01).

LD testing was also influenced by prior testing, significant effects of Rearing and Prior Testing, and interactions between Rearing and Sex, Rearing and Prior Testing, and Rearing, Sex, and Prior Testing on the latency to enter the dark zone in the LD test (Rearing: *F*_1,94_ = 9.06, *p* = 0.003, Sex: *F*_1,94_ = 19.5, *p* = 2.7 × 10^-5^, Testing: *F*_1,94_ = 21, *p* =1.4 × 10^-5^, Rearing × Sex: *F*_1,94_ = 7.26, *p* = 0.008, Rearing × Testing: *F*_1,94_ = 4.16, *p* = 0.044, Sex × Testing: *F*_1,94_ = 1.21, *p* = 0.28, Rearing × Sex × Testing: *F*_1,94_ = 5.54, *p* = 0.021) (Fig. 6B). MS male rats not previously tested on the LD test during the juvenile period took a significantly longer time to enter the dark zone than those tested during the juvenile period (Tukey’s *p* = 0.0043). This was also true for untested female rats regardless of Rearing condition, which took significantly less time to enter the dark zone (Tukey’s *p* = 0.0025).

## DISCUSSION

Here, we developed a severe form of MS in rat, consisting of 6-hour daily separation from P2 through P21 at unpredictable times, with the goal of detailed characterization of biobehavioral profiles across the lifespan following maximal disruption of normal dam-pup interactions. In this comprehensive study we assessed maternal anxiety-like behavior, pup maternal odor preference, juvenile play behavior, sociability, juvenile and adult anxiety-like behaviors, stress coping, and glucocorticoid stress responses. We also addressed sibling versus cross-housing conditions and the effects of repeated testing as potential confounders. Overall, our study showed minimal effects on long-term behavioral outcomes.

### MS increases preference for maternal odor in male rats

Our MS paradigm increased maternal odor preference in recently weaned MS male but not female rats. These results are consistent with a prior study showing the preference for odors associated with the dam to be enhanced by early life adversity (Raineki et al., 2010) and might be indicative of conditioned enhanced attachment to neglectful caregiving. Additionally, in a mouse model of MS, it was found that control mice prefer maternal odor initially (P10), but that this attraction begins to weaken as the mice age (P14). In contrast, this weakening did not occur in MS mice, which preferred their maternal odors both at P10 and at P14 (Thomas et al., 2010). However, neither of these studies differentiated between male and female behavior, making them difficult to compare to our sex-specific findings.

### MS does not influence social preference but does increase play behavior in juvenile females

Contrary to a number of reports showing decreased sociability following MS (Maciag, 2002; Tsuda et al., 2011; Farrell et al., 2016), social interaction testing during the juvenile and adult periods revealed no differences between groups in our model. All rats preferred to spend time in the chamber with an unfamiliar rat over a novel object, and this preference was strongest during the juvenile period.

However, a small and sex-specific difference in sociability was seen in juvenile play behavior, with MS females spending significantly more time playing than C female on the last day of testing (P38). Others who have tested juvenile play in MS-exposed rodents have found mixed results, with some groups finding no differences (Bodensteiner et al., 2014) and others finding an increase in aggressive play fighting (boxing) behaviors in female rats (Zimmerberg and Sageser, 2011). Although our MS female finding suggests enhanced propensity for social interaction, it might also reflect diminished ability to control social interactions. Interestingly, such an absence of social approach regulation has been seen in Romanian orphans (Kaler and Freeman, 1994; Chisholm, 1998).

### MS decreases anxiety-like behavior across the lifespan

Many studies have demonstrated that MS increases anxiety-like behavior as assessed in the open field and EPM tests in adulthood (Daniels et al., 2004; Cao et al., 2016; Shin et al., 2016; Bondar et al., 2018). Yet, negative (Millstein and Holmes, 2007), opposing (Leon Rodriguez and Duenas, 2013), or mixed effects (Markostamou et al., 2016; Dandi et al., 2018) have also been reported. Similarly, on the LD test, some found MS-induced deficits in juvenile rats (Chocyk et al., 2013) and adult rats (Wang et al., 2012), while others found a decrease in anxiety-like behavior in adult rats (Zhang et al., 2014) and still others found no effects (Tsuda and Ogawa, 2012). In our model, while absolute differences were modest, MS was overall associated with decreased anxiety-like behaviors on the LD test during both juvenile and adult periods. Housing condition played a role, with sibling-housing offering a small but significant additional protection against anxiety-like behavior in adult MS male rats.

### MS negatively impacts coping behaviors in MS male rats

Our only result consistent with our initial hypothesis was on the FST, where MS male rats showed increased despair behaviors (immobility) and decreased coping behaviors (climbing). Notably, MS-driven effects in males were unlikely to be related to weight differences as MS male rats weighed less than C male rats in adulthood, which would predict more ability to climb. It is well-established that early-life stressors, especially as modeled through MS, are a risk factor for developing depressive-like behaviors (Lee et al., 2007; Marais et al., 2008; Bian et al., 2015; Amiri et al., 2016; Masrour et al., 2018; Ruiz et al., 2018).

Parameters such as immobility are measures of depressive-like or coping behaviors and are sensitive to antidepressant treatment (Porsolt et al., 1977). Our results are in line with previous studies showing MS rats display decreased coping behaviors on the FST (Veenema et al., 2006; Desbonnet et al., 2010; Bian et al., 2015; Amiri et al., 2016; Genty et al., 2018; Masrour et al., 2018; Ruiz et al., 2018). Additionally, and somewhat unexpectedly (Kokras et al., 2015), our results also showed strong sexually dimorphic behavior on the FST, with females spending overall less time immobile and more time struggling compared to males, indicating better coping strategies.

### MS does not affects glucocorticoid responses to stress

As with most endpoints, MS has been associated with increased (Rincel et al., 2016; Ruiz et al., 2018), decreased (Biggio et al., 2018), or no effect (Rana et al., 2016) on corticosterone levels. When analyzing hypothalamic-pituitary-adrenal (HPA)-axis reactivity, the results have also been mixed, although a majority of studies have found increased reactivity (Lajud et al., 2012). In our study, there was no significant effect of MS on either basal or stress-induced serum corticosterone levels. However, as widely reported in the literature (Babb et al., 2013; Albrechet-Souza et al., 2020), we did observe a higher corticosterone peak response in female rats, correlating with the increased female struggling time observed in the FST.

### Conclusion

Our results further highlight the complex landscape of the rodent MS literature, which contains reports of both negative (Stanton et al., 1988; Kambali et al., 2019), positive (Zhang et al., 2014; Lundberg et al., 2017a) and null (Lehmann et al., 2000) effects of MS on future biobehavioral profiles. These conflicting findings are like due to the exact experimental design, including rodent species and strain, housing conditions, duration and exact time in the diurnal cycle for the separation, developmental period during which separation occurs, whether or not separation also extends to siblings, and post-weaning housing conditions.

It is interesting to note other studies employing prolonged MS paradigm from P1/P2 to P21 also have found minimal (Lundberg et al., 2017b) or positive behavioral effects (McIntosh et al., 1999), while most studies reporting negative outcomes use separation that ends around P14. The developing HPA axis in the rat undergoes a stereotypic process of a stress hyporesponsive period (SHRP) between P4 and P14, during which corticosterone responses are minimal or non-existent (Levine, 2001). Interestingly, MS-induced changes in dendritic spines in the anterior cingulate cortex, a key prefrontal region involved in emotional regulation, show distinct patterns depending on timing of the exposure, with decreased spine density in MS prior to SHRP, no changes in dendritic spines with MS during the SHRP and increases in dendritic spines with MS after the SHRP (Bock et al., 2005). Therefore, it is conceivable that our protocol led to initial deleterious neurocircuit changes during the first phase, with compensatory mechanisms developing during the later phase to ameliorate long-term maladaptive consequences.

## ACKNOWLEDGEMENTS

This work was sponsored by generous donations from Fleur Fairman, John and Rainy Erwin, and Einhorn Collaborative.

## Notes

### Competing Interest Statement

The authors have declared no competing interest.

